# *Adrβ2* in skeletal muscle cells is *required* for exercise-induced *Pgc1α* but not for metabolic benefits of exercise on diet-induced obesity

**DOI:** 10.64898/2026.03.27.714812

**Authors:** Marco Galvan, Mina Fujitani, Jasmine Dushime, Safia Baset, Bandy Chen, Shreya Thomas, Carlos M. Castorena, Joel K. Elmquist, Teppei Fujikawa

**Affiliations:** Center for Hypothalamic Research, Department of Internal Medicine, UT Southwestern Medical Center, Dallas, Texas, USA; Department of Neuroscience, UT Southwestern Medical Center, Dallas, Texas, USA; Department of Pharmacology, UT Southwestern Medical Center, Dallas, Texas, USA; Peter O’Donnell Jr. Brain Institute, UT Southwestern Medical Center, Dallas, Texas, USA; Institute of Human Life and Ecology, Osaka Metropolitan University, Osaka, Japan

**Keywords:** *Adrβ2*, exercise, skeletal muscle, metabolism, sympathetic nervous system

## Abstract

β2-Adrenergic receptor (*Adrβ2*) is the most abundant form of adrenergic receptors in skeletal muscle. Our previous studies have shown that the ventromedial hypothalamic nucleus (VMH) regulates metabolic benefits of exercise, potentially by skeletal muscle *Adrβ2*. Although a large body of literature has shown the importance of *Adrβ2* on skeletal muscle physiology, it remains unexplored whether skeletal muscle *Adrβ2* contributes to metabolic benefits of exercise, such as prevention of diet-induced obesity (DIO). Here, we generated mice lacking *Adrβ2* in skeletal muscle cells (SKM*^Adrβ2^*) and tested whether SKM*^Adrβ2^* is *required* for metabolic benefits of exercise on DIO. Deletion of SKM*^Adrβ2^* completely abolished the induction of *peroxisome proliferator-activated receptor gamma coactivator 1-alpha* (*Pgc-1α*) in skeletal muscle by β2-agonist, which is a potent activator of *Pgc-1α*. Exercise upregulates *Pgc-1α,* which regulates a broad range of skeletal muscle physiology, including hypertrophy and mitochondrial function. Deletion of SKM*^Adrβ2^* hampers augmented *Pgc-1α* in skeletal muscle by a single bout of exercise. Intriguingly, we found that deletion of SKM*^Adrβ2^* increased endurance capacity. Further, our data showed that body weight in DIO mice lacking SKM*^Adrβ2^* is comparable to that of control DIO mice during exercise training, suggesting that deletion of SKM*^Adrβ2^* did not affect the metabolic benefits of exercise in DIO. Collectively, our data indicate that SKM*^Adrβ2^* contributes to exercise-induced transcriptional changes and endurance capacity, however, it is not *required* for exercise benefits on bodyweight in DIO mice.

## Introduction

Exercise has a spectrum of metabolic benefits, yet mechanisms underlying metabolic benefits of exercise have remained unclear. In particular, neuronal and molecular mechanisms by which nervous systems contribute to metabolic benefits of exercise have just begun to be explored^1,2^. Several studies have shown that the central nervous system (CNS) contributes to the metabolic benefits of exercise via the sympathetic nervous system (SNS)^2–4^. Within the CNS, hypothalamic neurons are key to metabolic benefits of exercise^2^. For instance, microinjection of β-blockers into the ventromedial hypothalamic nucleus (VMH), which regulates the SNS, led to alterations in energy substrate utilization during exercise^5–7^. Steroidogenic factor-1 (SF-1) neurons in the VMH are key to metabolic benefits of exercise on high-fat diet (HFD)-induced obesity^1^, and they regulate transcriptional events in skeletal muscle via the SNS^8,9^. Further, exercise activates some population of SF-1 neurons, and neuronal activation of SF-1 neurons can enhance endurance capacity^10^. Collectively, studies have shown that the CNS-SNS axis plays a significant role in the regulation of skeletal muscle physiology and metabolism in the context of exercise.

The SNS utilizes epinephrine (Epi) and norepinephrine (NE) as neurotransmitters to communicate with peripheral tissues^11^. Among nine adrenergic receptors for Epi and NE^12^, β2-Adrenergic receptor (*Adrβ2*) is the most abundant form of adrenergic receptors in skeletal muscle^13^. A large body of literature has shown the role of *Adrβ2* in skeletal muscle function and metabolism^13^. For instance, injections of agonists of *Adrβ2* (β2-agonist), lead to drastic transcriptional changes^14,15^, ultimately increasing skeletal muscle weight and function^16–18^. Further, β2-agonist administrations improve insulin sensitivity and overall glucose metabolism^19–21^, and *Adrβ2* in skeletal muscle cells is required for this action^15^. These metabolic actions of skeletal muscle *Adrβ2* likely contribute to beneficial effects of exercise on metabolism driven by the CNS-SNS axis. However, there are no concrete studies investigating the role of skeletal muscle *Adrβ2* on metabolic actions in the context of exercise. Here, we used genetically-engineered mice that lack *Adrβ2* in skeletal muscle cells (SKM*^Adrβ2^*) and investigated whether SKM*^Adrβ2^* is required for metabolic benefits of exercise.

## Results

### Generation of mice lacking *Adrβ2* in skeletal muscle cells

To generate mice lacking *Adrβ2* in skeletal muscle cells, we crossed HAS-MCM (RRID:IMSR_JAX:025750)^22^ and *Abrb2* floxed mice^23^ (ΔSKM*^Adrβ2^*). HAS-MCM mice express tamoxifen-inducible Cre recombinase, thus *Abrb2* in skeletal muscle cell of ΔSKM*^Adrβ2^* would be knocked down upon tamoxifen injection. Studies have shown that i.p. injection of tamoxifen has more drastic impacts on metabolism compared to other routes such as oral intake of tamoxifen^24^. Thus, we used tamoxifen-diet (Tam-diet) to induce Cre activity in skeletal muscle cells. Food intake decreased during the first week of the Tam-diet but was restored from the second week (**Fig. 1A**). Along with decreased food intake, during tamoxifen-diet feeding, body weight was decreased to approximately 5-10% from the prior body weight (BW), but reduced BW was reversed quickly within one week after switching to normal diet (**Fig. 1B**). Tam-diet significantly reduced *Adrβ2* mRNA in skeletal muscle (**Fig. 1C**), but not smooth muscle or cardiac muscle (**Fig. 1D and E**), suggesting that deletion of *Adrβ2* occurs specifically in skeletal muscle cells. In all experiments outlined from here, mice had at least a four-week recovery time from the Tam-diet.

**Figure 1.**
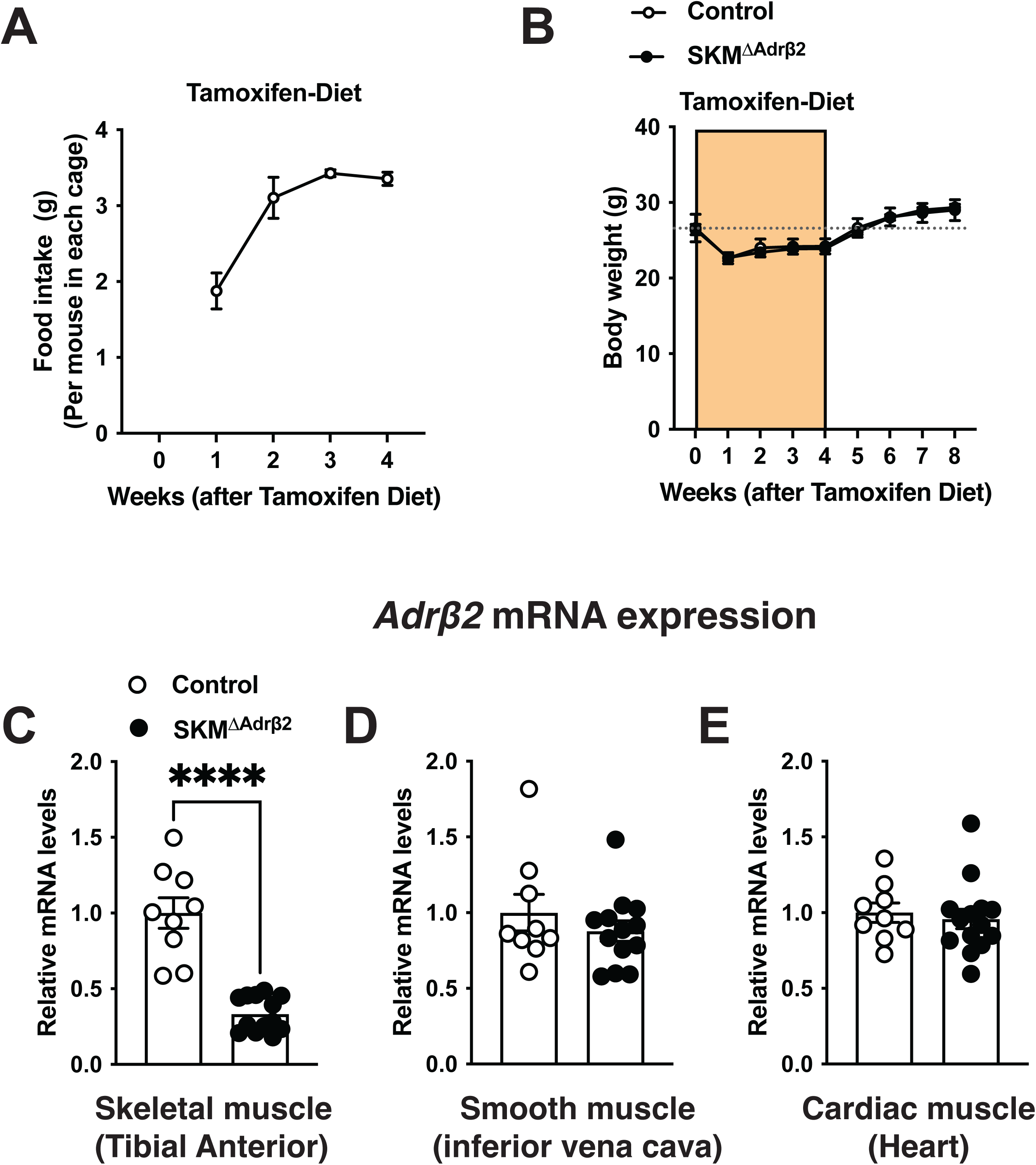
Adult ablation of *Adrβ2* in skeletal muscle by tamoxifen diet. (**A**) Food intake during tamoxifen diet (TAM-diet) feeding. The food weight was measured as the total weight of the cage and divided by the number of mice. (**B**) The body weight curve during and after TAM-diet feeding. The orange color indicates the time-window of TAM-diet feeding. mRNA levels of *Adrβ2* in (**C**) tibial anterior skeletal muscle, (**D**) inferior vena cava, and (**E**) heart in mice lacking *Adrβ2* in skeletal muscle cells and littermate wild-type control. Values are mean ± S.E.M. ****p < 0.0001. A detailed statistical analysis is described in Supplemental Table 3.

### *Adrβ2* in skeletal muscle cells is required for exercise-induced *Pgc-1α* mRNA

We investigated whether ΔSKM*^Adrβ2^* disrupts *Adrβ2* function in skeletal muscle. To test this, we injected β2-agonist, clenbuterol, into mice and examined mRNA levels of *peroxisome proliferator-activated receptor gamma coactivator 1-alpha* (*Pgc-1α*) in skeletal muscle (**Fig. 2A and B**). β2-agonist is one of the most potent drugs that can increase *Pgc-1α* in skeletal muscle^25–27^. We found that ΔSKM*^Adrβ2^* completely blocked the induction of *Pgc-1α* mRNA by β2-agonist (**Fig. 2C**), indicating ΔSKM*^Adrβ2^* diminishes *Adrβ2* function in skeletal muscle, as well as *Pgc-1α* induction by β2-agonist solely relies on *Adrβ2* in skeletal muscle cells, but not other cell types, such as smooth muscle and satellite cells in skeletal muscle. Then, we asked whether SKM*^Adrβ2^* is required for exercise-induced *Pgc-1α* mRNA in skeletal muscle (**Fig. 3A and B**). *Pgc-1α* regulates mitochondrial function, glucose and lipid metabolism, and growth of skeletal muscle^28–30^, and exercise dramatically increases *Pgc-1α* mRNA in skeletal muscle^1,26,31^. As expected, after a single exercise, *Pgc-1α* mRNA in skeletal muscle, in particular isoform 2-4, was increased in control compared to sedentary groups (**Fig. 3C**). However, exercised ΔSKM*^Adrβ2^* mice did not show significant increases in *Pgc-1α* mRNA in skeletal muscle compared to sedentary ΔSKM*^Adrβ2^* mice, suggesting that SKM*^Adrβ2^* is required for exercise-induced *Pgc-1α* mRNA (**Fig. 3C)**.

**Figure 2.**
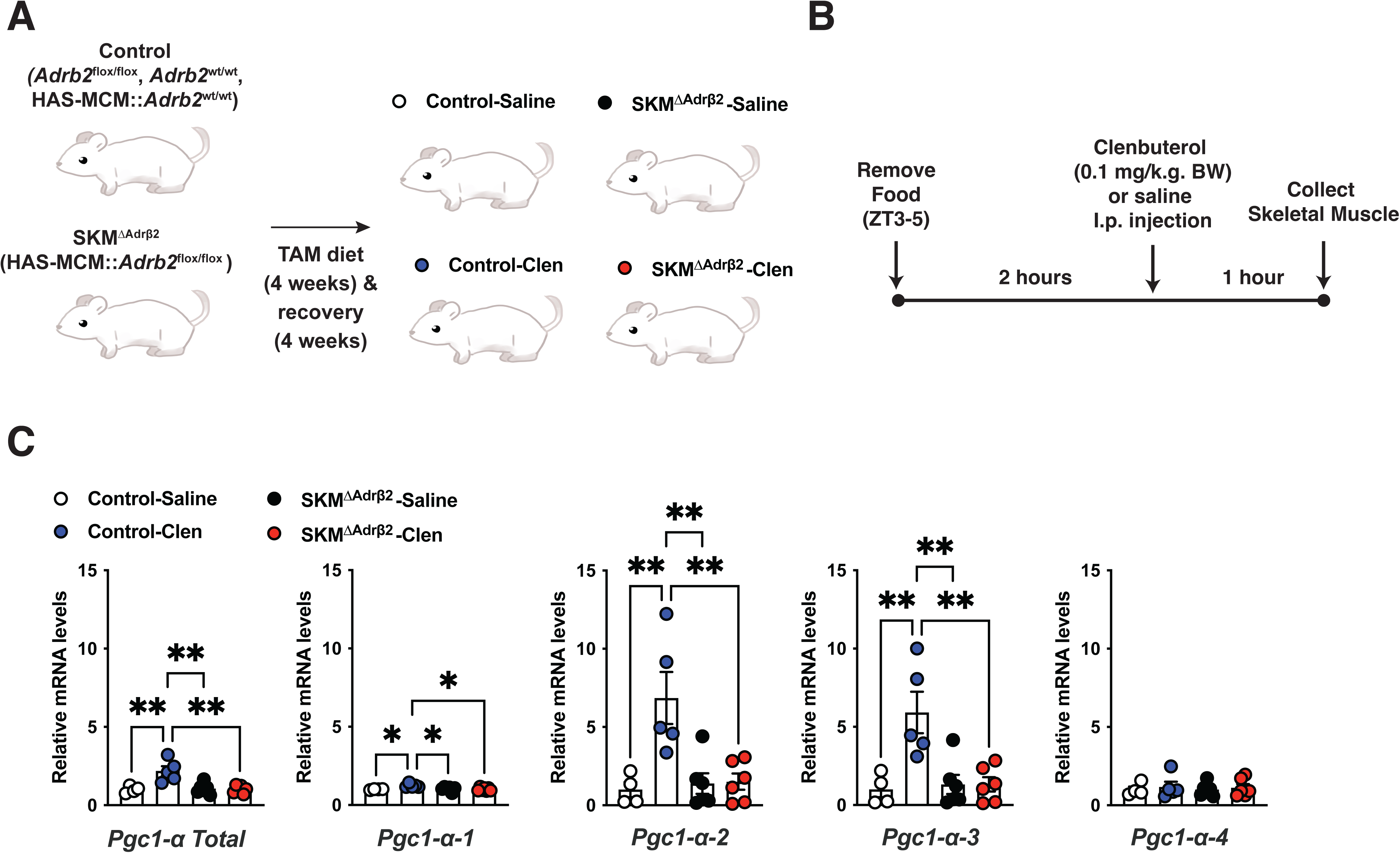
*Adrβ2* in skeletal muscle cells is *required* for β2agonist-induced *Pgc-1α* mRNA in skeletal muscle. (**A**) Schematic design of breeding and tamoxifen strategy. (**B**) Experimental design. (**C**) mRNA levels of *Pgc1-α* isoforms in TA muscle of ΔSKM*^Adrβ2^* mice after clenbuterol injection. Values are mean ± S.E.M. **p <0.01, *p < 0.05. A detailed statistical analysis is described in Supplemental Table 3.

**Figure 3.**
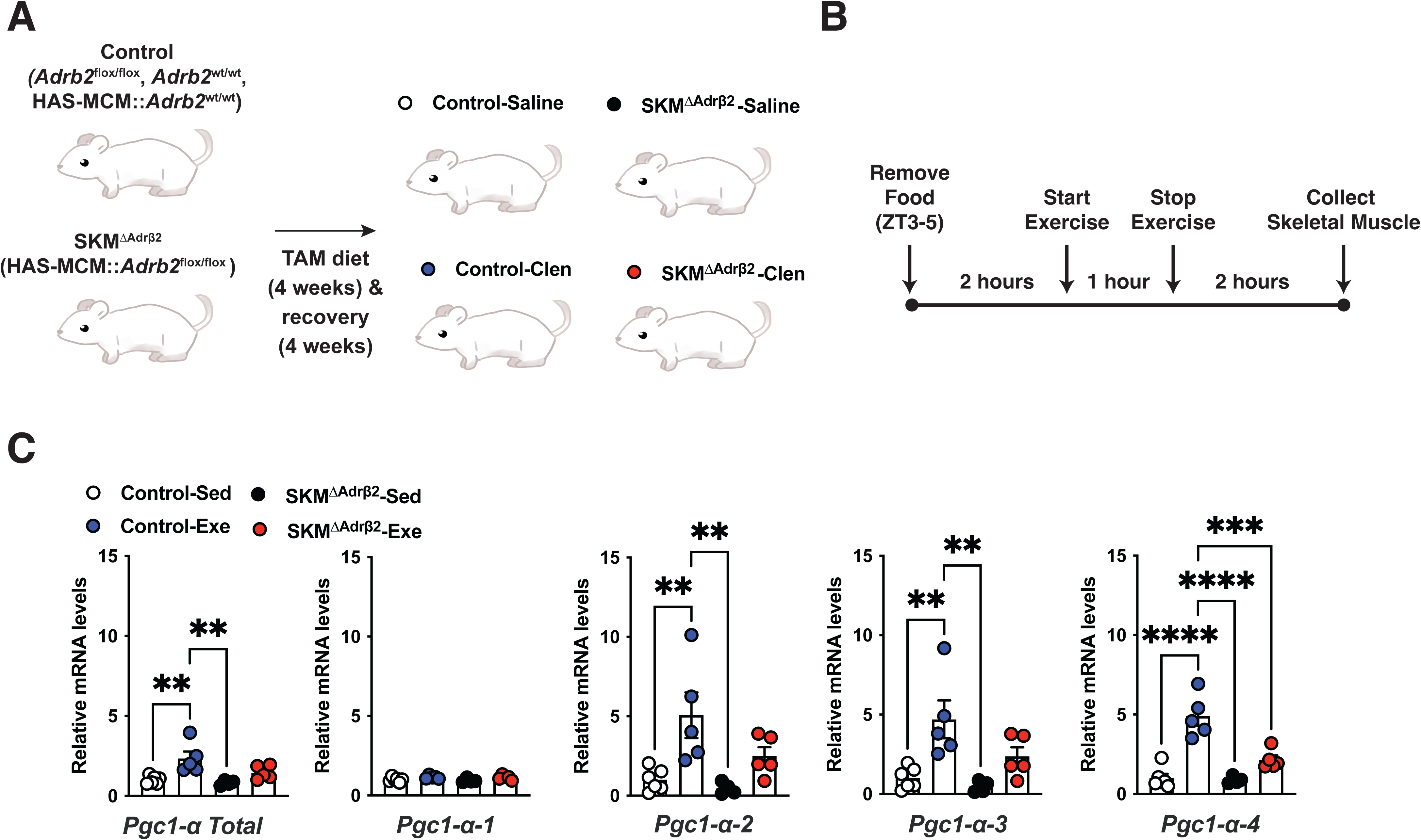
*Adrβ2* in skeletal muscle cells is required for exercise-induced *Pgc-1α* mRNA in skeletal muscle. (**A**) Schematic design of breeding and tamoxifen strategy. (**B**) Experimental design. (**C**) mRNA levels of *Pgc1-α* isoforms in TA muscle of ΔSKM*^Adrβ2^* mice after exercise. Values are mean ± S.E.M. **** p < 0.0001, *** p < 0.001, ** p <0.01. A detailed statistical analysis is described in Supplemental Table 3.

### Deletion of *Adrβ2* in skeletal muscle cells increases endurance capacity

The original study that generated whole-body *Adrβ2* knockout mice (*Adrβ2*^KO^) demonstrated that *Adrβ2*^KO^ mice exhibit increased endurance capacity^32^. We investigated whether the improved endurance capacity in *Adrβ2*^KO^ mice results from SKM*^Adrβ2^*. To do so, we tested the endurance capacity of ΔSKM*^Adrβ2^* mice by a progression endurance capacity test that we used before^1^. Intriguingly, ΔSKM*^Adrβ2^* mice ran significantly longer than control mice (**Fig. 4A**), suggesting that SKM*^Adrβ2^* controls endurance capacity in mice. There were no significant differences in blood and lactate levels immediately after the endurance test between groups (**Fig. 4B and C**). The previous study showed that *Adrβ2*^KO^ mice do not exhibit increased oxygen consumption but instead have increased fat oxidation as indicated by a lower respiratory exchange ration^32^. Collectively, these data suggest that SKM*^Adrβ2^* regulates the use of carbohydrate/fat fuels during endurance exercise.

**Figure 4.**
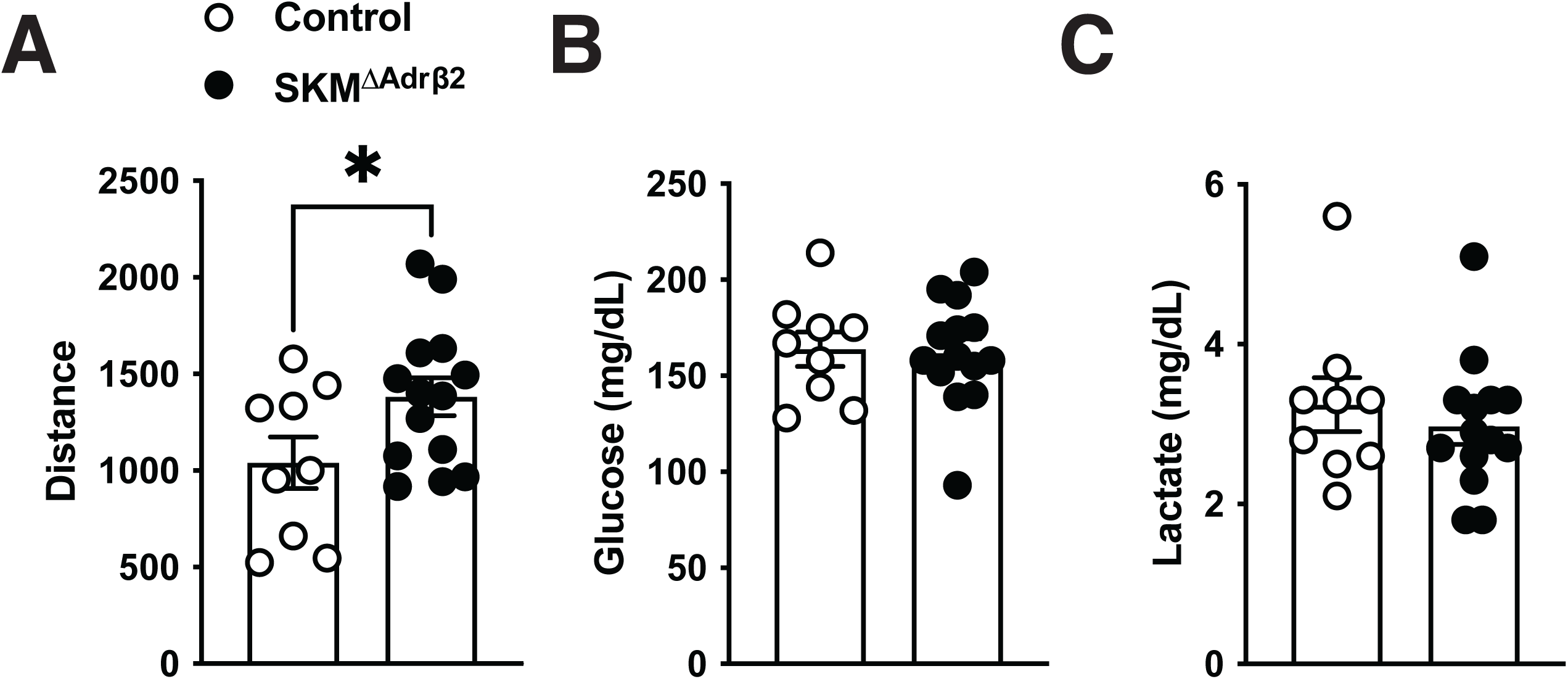
Adult ablation of *Adrβ2* in skeletal muscle extends endurance capacity. (**A**) Endurance capacity, (**B**) blood glucose, and (**C**) lactate levels immediately after the endurance capacity test in ΔSKM*^Adrβ2^* mice. Values are mean ± S.E.M. *p < 0.05. A detailed statistical analysis is described in Supplemental Table 3.

### Deletion of *Adrβ2* in skeletal muscle cells does not affect metabolic benefits of exercise on high-fat diet-induced obesity

Finally, we tested whether SKM*^Adrβ2^* is required for beneficial effects of exercise on HFD-induced obesity. In mice, exercise effectively prevents HFD-induced obesity even middle intensity of exercise, which we used in this study^1^. Of note, high intensity and prolonged exercise training are required to achieve weight loss in humans if there is no food intake intervention^33–36^. In the first week, all mice were fed HFD, which 60% of calories are from fat, and simultaneously acclimated to the treadmill apparatus. Mice were trained for 4 weeks along with HFD feeding. As expected, exercise training significantly suppressed typical body weight gain resulting from HFD feeding as observed in the control group (**Fig. 5**). Intriguingly, ΔSKM*^Adrβ2^* did not affect exercise-induced prevention of gained weight in mice (**Fig. 5**). These results suggest that expression of SKM*^Adrβ2^* is not necessary for beneficial effects of exercise to prevent HFD-induced obesity.

**Figure 5.**
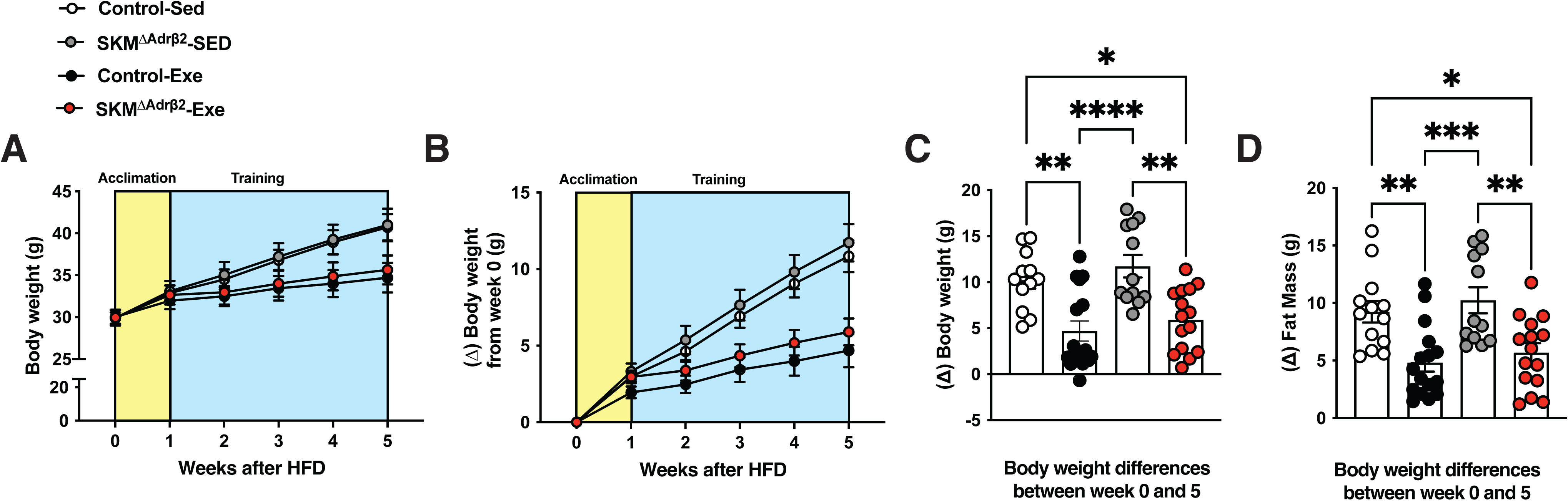
*Adrβ2* in skeletal muscle cells is not *required* for metabolic benefits of exercise on high-diet induced obesity. (**A**) Time course of body weight, and (**B**) time course of differences in body weight prior to exercise training in ΔSKM*^Adrβ2^* mice. (**C**) body weight, and (**D**) fat mass differences from the baseline after 5 weeks exercise training in ΔSKM*^Adrβ2^* mice. Values are mean ± S.E.M. **** p < 0.0001, *** p <0.001, **p < 0.01, *p < 0.05. A detailed statistical analysis is described in Supplemental Table 3.

## Discussion

In this study, we demonstrated that SKM*^Adrβ2^* is *required* for exercise-induced *Pgc-1α* mRNA in skeletal muscle, yet is not required for beneficial effects of exercise on HFD-induced obesity. Human genome wide association studies (GWAS) have shown that *Adrβ2* is highly correlated with obesity, diabetes, and other metabolic diseases^37–39^, highlighting the importance of *Adrβ2* in metabolic homeostasis in humans. *Adrβ2* is the most abundant form of adrenergic receptors in skeletal muscle in human and rodents like mice, although it is expressed in other metabolically important organs including the liver, brain, and pancreas^40^. *Adrβ2* in skeletal muscle is key to many metabolic processes in skeletal muscle. β2-agonist administrations improve insulin sensitivity and overall glucose metabolism^19–21^, and *Adrβ2* in skeletal muscle cells is required for this action^15^. We did not assess whether SKM*^Adrβ2^* contributes to other metabolic benefits of exercise. SKM*^Adrβ2^* mediates pharmacological improvements of glucose metabolism by β2-agonist^15^. Also, it seems that SKM*^Adrβ2^* is key to maintaining glucose homeostasis in young^15,19,21^ and potentially age^39,41^. Thus, it would be intriguing to test whether SKM*^Adrβ2^* contributes to improvements in glucose metabolism by exercise training in both young and aged subjects. Future studies will be warranted to reveal the role of SKM*^Adrβ2^* in the regulation of glucose metabolism in the context of exercise.

Our previous studies have shown that VMH neurons expressing SF-1 (VMH^SF-1^) contribute to metabolic benefits of exercise including increased energy expenditure, reduced body weight when HFD is fed, and augmented skeletal muscle mass^1^. Knockdown of SF-1 in the VMH hampers the beneficial effects of exercise on HFD-induced obesity^1^. Activation of VMH^SF-1^ neurons increases *Pgc-1α* mRNA in skeletal muscle^8^, and the action is mediated by SKM*^Adrβ2^* ^9^. Several studies have shown that the sympathetic nervous system, including *Adrβ2* actions, mediates VMH-regulation of peripheral metabolism^42–44^. Thus, we postulated that SKM*^Adrβ2^* is key to metabolic benefits of exercise on HFD-induced obesity that is mediated by the CNS as VMH neurons. However, our current study indicates that SKM*^Adrβ2^* does not play a significant role in VMH-regulation of body weight in the context of exercise. Several mechanisms may contribute to VMH regulation of body weight in the context of exercise. For instance, the VMH regulates adipose tissue biology via the SNS^42,45–49^. The activation of the VMH increases glucose uptake into brown adipose tissues (BAT), resulting in increases in BAT temperature^49^. Further, optogenetic activation of cells expressing *cckbr* neurons in the VMH increases lipolysis, and these actions are blocked by β3-blocker injections^50^. Therefore, instead of adrenergic receptors in skeletal muscle, adrenergic receptors in adipose tissue may contribute to VMH regulation of body weight in the context of exercise.

ΔSKM*^Adrβ2^* mice exhibit a longer endurance capacity compared to control mice (**Fig 4**), which is a comparable phenotype to *Adrβ2*^KO^ mice^32^. Clenbuterol injections increase glucose utilization and oxidation, which are mediated by SKM*^Adrβ2^* ^15^. Mechanistically, clenbuterol injections reduce genes related to fatty acid uptake and oxidation in skeletal muscle^15^. Intriguingly, *Adrβ2*^KO^ mice exhibit reduced glucose utilization during the endurance test^32^. Collectively, these data indicate that SKM*^Adrβ2^* promotes a shift in energy utilization from fat to glucose during exercise. The greater endurance observed in ΔSKM*^Adrβ2^* mice likely results from increased fatty acid oxidation during the initial phase of exercise^32^, which can preserve glucose resources such as glycogen. The availability of a carbohydrate substrate is a critical factor in the progressive endurance test. Animals rely on the glycolytic pathway to produce ATP at higher exercise intensities, which require greater oxygen consumption^51^. Further studies are required for the molecular mechanisms underlying the regulation of energy substrate partitioning by SKM*^Adrβ2^*.

## Conclusion

Although SKM*^Adrβ2^* is well recognized as a key molecule to regulate skeletal muscle metabolism, its contributions of SKM*^Adrβ2^* to beneficial effects of exercise on metabolism remain unclear. We found that ΔSKM*^Adrβ2^* extends the endurance capacity but does not affect metabolic benefits of exercise on HFD-induced obesity in mice. There are several limitations in the current study. We only investigate male mice. The SNS is well known to regulate physiology in a sexual dimorphism manner^52,53^. Future studies need to address this point. Although *Adrβ2* is the primary form of adrenergic receptors in skeletal muscle, the expression pattern of other adrenergic receptors in skeletal muscle or non-skeletal muscle tissue differs^40^, which may ultimately affect *Adrβ2* actions or compensate for the lack of *Adrβ2* in skeletal muscle. For instance, the most abundant adrenergic receptor in murine adipose tissue is *Adr*β*3*^40^. However, human adipose tissue barely expresses *Adr*β*3* and the primary adrenergic receptor is *Adrβ2*^54^. Therefore, the current findings using mice may not translate directly into human physiology. Finally, the metabolic benefits of exercise are widespread across many physiological processes. In addition, the benefits depend on the exercise training regimen, such as resistance training vs. endurance training, the intensity of training, and the length of training^28,29,55,56^. We only investigated the metabolic benefits on HFD-induced obesity in mice. Therefore, future studies are warranted to further investigate the role of SKM*^Adrβ2^* in various settings of exercise training on other metabolic benefits of exercise, such as glucose metabolism.

## Materials and Methods

### Genetically-engineered mice and mouse husbandry

HAS-MCM (RRID:IMSR_JAX:025750)^22^ were purchased from the Jackson Laboratory. *Abrb2* floxed mice^23^ were obtained from Dr. Florent Elefteriou at Baylor College of Medicine with the permission of Dr. Gerard Karsenty at Columbia University. Ear or tail gDNA was collected from each mouse to determine its genotype. A KAPA mouse genotype kit was used for PCR genotyping (Roche, US). The sequences of genotyping primers and expected band sizes for each allele are described in **Supplemental Table 1**. Mice were housed at room temperature (22–24 C) with a 12 hr light/dark cycle (lights on at 7 AM and 6 AM during daylight saving time, respectively) and fed a normal chow diet (2016 Teklad global 16% protein rodent diets, Envigo, US). 60% kCal fat diet (Research Diets Inc., D12492; 5.21Kcal/g) was used as a high-fat diet. To induce Cre activity, mice were fed with Tamoxifen diet (Envigo, US; Catalog number TD.130859) for 4 weeks. Magnetic-resonance whole-body composition analyzer (EchoMRI, US) was used to analyze the fat and lean mass of mice. All procedures using mice were approved by the Institutional Animal Care and Use Committee of UTSW, and animal experiments were performed at UTSW.

### Clenbuterol injection

Clenbuterol hydrochloride (Sigma Aldrich, US. Catalog# 1134674) was dissolved in sterile saline at a concentration of 5 mg/mL as the stock solution. The stock solution was stored at −80 °C. On the day of the experiment, the stock solution was diluted to 0.01 mg/mL and i.p. injected into the mice (0.1 mg/kg body weight; e.g., 30 grams of mice received 300 µL of diluted solution). Tissues were collected into the tissue collection tubes 1 hour after injection, frozen immediately using dry ice, and kept at −80 °C.

### Exercise

We used the exercise protocol which was described in our previous study^1^. A 10° treadmill incline was used in all experiments. An electrical shock (0.25 mA at 163 V and 1 Hz) was used to encourage mice to run. At ZT3-5, food was removed from both exercise and sedentary groups. Two hours after food removal, treadmill running session was carried out. A water bottle was removed from sedentary groups during the exercise session. Food and water were returned to cages immediately following the exercise session unless otherwise stated.

For a one-time exercise session, on day 1, all mice were acclimated to the treadmill apparatus at a speed of 8 m/min for 5 minutes. On day 2, all mice ran at 8 m/min for 5 minutes followed by 10 m/min for 10 minutes. Acclimated mice were then rested in their home cages on day 3 and 4. Mice were randomly assigned to exercise or sedentary groups at this point. On day 5, exercise group run at 15 m/min for 60 min. Two hours after exercise, tissues were collected as outlined above.

For an endurance capacity test, a progressive running paradigm was used^1^. Mice were acclimated in the same manner as a one-time exercise session. On day 5, mice began the exercise training at a speed of 10 m/min for 40 minutes. After the initial 40 minutes, the speed was increased at a rate of 1 m/min every 10 minutes until the speed reached 13 m/min; at which point, the speed was increased at a rate of 1 m/min every 5 minutes until all mice were exhausted. Time to exhaustion was defined as the point mice spent more than 5 seconds on the electrical shocker.

For prolonged training, mice were acclimated starting at week 0 as described previously^1^. Mice were either maintained on HFD or chow from the week 0. During week 1 of training, the speed was gradually increased day by day until the speed reached 15 m/min. From the week 2 onwards, mice ran at a speed of 10 m/min for 5 minutes and the speed was increased at a rate of 1 m/min every 1 minute until the speed reached 15 m/min. The speed was maintained for 51 minutes (total run time is 60 minutes). The training bouts were performed 5 days/week (Monday-Friday).

### Assessment of mRNA

mRNA levels in tissues were determined as previously described ^1,57^. The sequences of primers for qPCR using SYBR are described previously ^31^ and in **Supplemental Table 2**.

### Graphic software, Data analysis, and statistical design

The data are represented as means ± S.E.M. GraphPad PRISM version 10 (GraphPad, US) was used for the statistical analyses and *P*<0.05 was considered as a statistically significant difference. A detailed analysis of each figure was described in **Supplemental Table 3**. The sample size was decided based on previous publications ^1,7,57–63^, and no power analysis was used. Figures were generated using PRISM version 10, Illustrator 2025, and Photoshop 2025 (Adobe Inc, US).

## Acknowledgment

We would like to thank Dr. Florent Elefteriou at Baylor College of Medicine and Dr. Gerard Karsenty at Columbia University for sharing *Abrb2* floxed mice. This study was supported by the National Institute of Health (To J.K.E., P01DK119130, R01DK088423, and R01DK100659, UTSW NORC P30DK127984), and by JSPS KAKENHI Grant Number 24K23124 to T.F.

## Contribution

M.G., M.F., and C.C. designed, performed, and analyzed experiments, and edited the manuscript. M.G. and M.F. equally contributed to this study, and they have the right to list themselves as the first author. J.D., S.B., B.C., and S.T., performed experiments. J.K.E. supervised and edited the manuscript. T.F. designed, performed, supervised, and analyzed experiments, and wrote and finalized the manuscript.

